# Multidimensional plasticity jointly contributes to rapid acclimation to environmental challenges during biological invasions

**DOI:** 10.1101/2022.06.16.496377

**Authors:** Xuena Huang, Hanxi Li, Noa Shenkar, Aibin Zhan

**Affiliations:** Research Center for Eco-Environmental Sciences, Chinese Academy of Sciences, 18 Shuangqing Road, Haidian District, Beijing 100085, China; University of Chinese Academy of Sciences, Chinese Academy of Sciences, 19A Yuquan Road, Shijingshan District, Beijing 100049, China; School of Zoology, George S. Wise Faculty of Life Sciences, Tel-Aviv University, 6997801 Tel-Aviv, Israel; The Steinhardt Museum of Natural History, Israel National Center for Biodiversity Studies, Tel Aviv University, Tel-Aviv, Israel

**Keywords:** Phenotypic plasticity, Biological invasion, Gene expression, Alternative splicing, Alternative polyadenylation

## Abstract

Rapid plastic response to environmental changes, which involves extremely complex underlying mechanisms, is crucial for organismal survival during many ecological and evolutionary processes such as those in global change and biological invasions. Gene expression is among the most studied molecular plasticity, while co- or post-transcriptional mechanisms are still largely unexplored. Using a model invasive ascidian *Ciona savignyi*, we studied multidimensional short-term plasticity in response to hyper- and hypo-salinity stresses, covering the physiological adjustment, gene expression, alternative splicing (AS), and alternative polyadenylation (APA) regulations. Our results demonstrated that rapid plastic response varied with environmental context, timescales, and molecular regulatory levels. Gene expression, AS, and APA regulations independently acted on different gene sets and corresponding biological functions, highlighting their non-redundant roles in rapid environmental adaptation. Stress-induced gene expression changes illustrated the use of a strategy of accumulating free amino acid under high salinity and losing/reducing them during low salinity to maintain the osmotic homoeostasis. Genes with more exons were inclined to use AS regulations, and isoform switches in functional genes such as *SLC2a5* and *Cyb5r3* resulted in enhanced transporting activities by upregulating the isoforms with more transmembrane regions. The extensive 3’-untranslated region (3’UTR) shortening through APA was induced by both salinity stresses, and APA regulation predominated transcriptomic changes at some stages of stress response. The findings here provide evidence for complex plastic mechanisms to environmental changes, and thereby highlight the importance of systemically integrating different levels of regulation mechanisms in studying initial plasticity in evolutionary trajectories.

## INTRODUCTION

In the past several decades, biogeographical patterns of many species have been rapidly reshaped globally with the intensification of human activities (Sandoval-Castillo et al. 2020; Wiens 2016), posing a challenge to the current survival of organisms. Rapid plastic response plays an important role in organismal survival and population persistence in adverse environments (Sandoval-Castillo et al. 2020). However, highly dynamic and complex plastic response has been reported when taking different plastic traits of interest, environmental contexts, and time scales into consideration (Fox et al. 2019; Metzger and Schulte 2018; Huang and Zhan 2021). Thus, untangling the complexity of plastic mechanisms during the response to environmental changes is crucial for understanding the organismal performance in many ecological and evolutionary scenarios, such as those in global change and biological invasions.

Biological invasions provide contemporary “natural experiments” for studying complex mechanisms of plastic response to environmental changes (Santi et al. 2020). During invasions, range expansions can occur at extremely large geographical scales, rapidly spanning distinct environmental regimes over relative short timescales (Zhan et al., 2010, 2015). When invasive species are initially introduced to novel environments, rapid plastic response is firstly stimulated to mitigate environmental stresses, and subsequently might participate in adaptive evolution after successful establishment. For example, the plastic response of producing more clonal propagules to water availability in the invasive sunflower *Helianthus tuberosus* facilitated the plant survival and persistence when introduced to riparian habitats (Bock et al. 2018). Marine invasive species are often introduced into novel habitats by human activity-mediated vectors such as ballast water and hull fouling during shipping (Ruiz and Carlton 2003). During the transport stage of transoceanic shipping, the drastic salinity shifts can be ∼ 15 ‰ in ballast tanks during a several-day voyage (Briski et al. 2013), and such fluctuation ranges can be even larger as fouling communities transported among ports (Bereza and Shenkar 2022), thus posing huge osmotic challenges to marine invasive species. The rapid plastic response of maintaining osmotic homoeostasis is crucial for the survival of marine invaders and subsequent successful invasions to environmentally distinct habitats (Jeffries et al. 2019; Posavi et al. 2020). However, most studies often focused on single regulation levels (e.g., Huang et al. 2017; Li et al. 2020; Chen et al. 2022), ignoring the intrinsic integrity and complexity of plastic response mechanisms.

Gene expression is among the most well-known molecular plastic traits, mainly because transcriptional regulations, which link genotypes and phenotypes, can be immediately induced to cope with environmental stresses during many ecological and evolutionary processes (Fu et al. 2021; Josephs 2021). Many studies have examined the roles of gene expression plasticity in organismal acclimation and adaptation to various environmental changes. For instance, the higher gene expression plasticity under acute thermal stresses was associated with enhanced thermal tolerance of a highly invasive inland siversides (*Menidia beryllina*) when compared to endangered native delta smelt (*Hypomesus transpacificus*), potentially conferring high invasiveness of silversides (Komoroske et al. 2021). Gene expression change, however, is not the only intermediate regulatory process linking genotypes and phenotypic traits. Alternative splicing (AS), as an important co- or post-transcriptional mechanism of generating multiple mRNA isoforms from a single gene, has received relatively less attention despite its importance in adaptation (Salisbury et al. 2021; Verta and Jacobs 2022). Several recent studies have demonstrated the contribution of AS to phenotypic novelty such as the caste differentiation of bumble bee (Price et al. 2018) and parallel ecological adaptation of ecotype-specific feeding morphology in a salmonid fish (Jacobs and Elmer 2021). Furthermore, genome-wide AS patterns could be rapidly reprogrammed at a short timescale to cope with environmental changes (Shalgi et al. 2014; Suresh et al. 2020; Tian et al. 2020), making AS a good candidate molecular plastic trait for studying initial plasticity. In addition to AS, alternative polyadenylation (APA) is another post-transcriptional mechanism of generating multiple transcript isoforms with different length of 3’-untranslated regions (3’UTR) from a single gene, mainly through differential usage of polyadenylation sites (PAS). APA isoforms with different 3’UTR length can present quite different mRNA characteristics such as distinct mRNA stability, cellular localization, and even coding sequences (Sadek et al. 2019), resulting in protein diversity or different phenotypic outcomes. Although accumulating evidence illustrates that APA should be involved in various biological processes (Zheng et al. 2018; Pereira-Castro and Moreira 2021), it remains largely unexplored and even overlooked when investigating the mechanisms of rapid response to environmental stresses.

All abovementioned molecular plastic changes, including gene expression, AS, and APA isoform switch, integrally affect the quantity and quality of mRNA and finally contribute to phenotypic variation in changing environments. It has been proposed that different types of plasticity should act on distinct environmental changes and different timescales (Fox et al. 2019; Metzger and Schulte 2018). Therefore, in order to untangle the complexity of initial plastic changes, it is needed to investigate the regulatory interplay among these mechanisms, as well as their relative importance during response to different environmental challenges. As such, we used a model marine invasive species, the solitary ascidian *Ciona savignyi*, to study its initial plastic response mechanisms to salinity changes during invasions. As one of the major bio-fouling animals, *C. savignyi* could colonize a variety of artificial facilities such as shipping vessels in an extremely high density to spread globally (Zhan et al. 2015). *Ciona savignyi* is usually considered to be a northern Asian native, but now has been widely recorded globally such as the coasts of North America and even New Zealand and Argentina in the South Hemisphere (Fofonoff et al. 2018; Smith et al. 2010; Zhan et al. 2015). During the transport stage, ascidian individuals can encounter drastic salinity shifts, and studies showed that they can survive a wide range of salinity from 18 ‰ to 40 ‰ (Fofonoff et al. 2018; Lambert and Lambert 2003). As an osmoconformer animal, the osmolarity of extra- and intra-cellular fluids can change with the external environments (Sokolov and Sokolova 2019), but the rapid regulation process has yet to be fully determined.

In order to systematically elucidate the underlying plastic mechanisms of salinity stress response, we used *C. savignyi* to conduct time-course hyper- and hypo-salinity challenges to investigate the enzyme activity, gene expression, AS, and APA plasticity. We aim to study the interplay of different plastic mechanisms at transcriptional and post-transcriptional regulatory levels and further identify key candidate genes and functional pathways responsible for rapid salinity acclimation.

## RESULTS

### Physiological response to salinity stresses

We detected significant and dynamic changes of four physiological indicators after salinity challenges. Specifically, the content of malondialdehyde (MDA) was rapidly accumulated soon after 1-hour of high salinity (HS) and low salinity (LS) challenge, maintained at a significantly high level at 24-hour of HS but significantly decreased at 24-hour of LS, and finally returned to the control level at 48-hour after HS and LS (Figure 1A). As for the antioxidant enzymes, the catalase (CAT) activity significantly decreased throughout the whole stress treatment process (Figure 1B), while the superoside dismutase (SOD) activity was significantly induced at 1-hour of HS and LS, remained rising at 24-hour of HS, but shifted to a repressed status after 48-hour of HS and LS (Figure 1C). The Na^+^/K^+^ ATPase activity was significantly induced across all three time-points under LS, but did not increase until 48-hour after HS (Figure 1D). These results illustrated that the antioxidant system and ion transport regulation were actively involved in the physiological response to salinity stresses.

**Figure 1.**
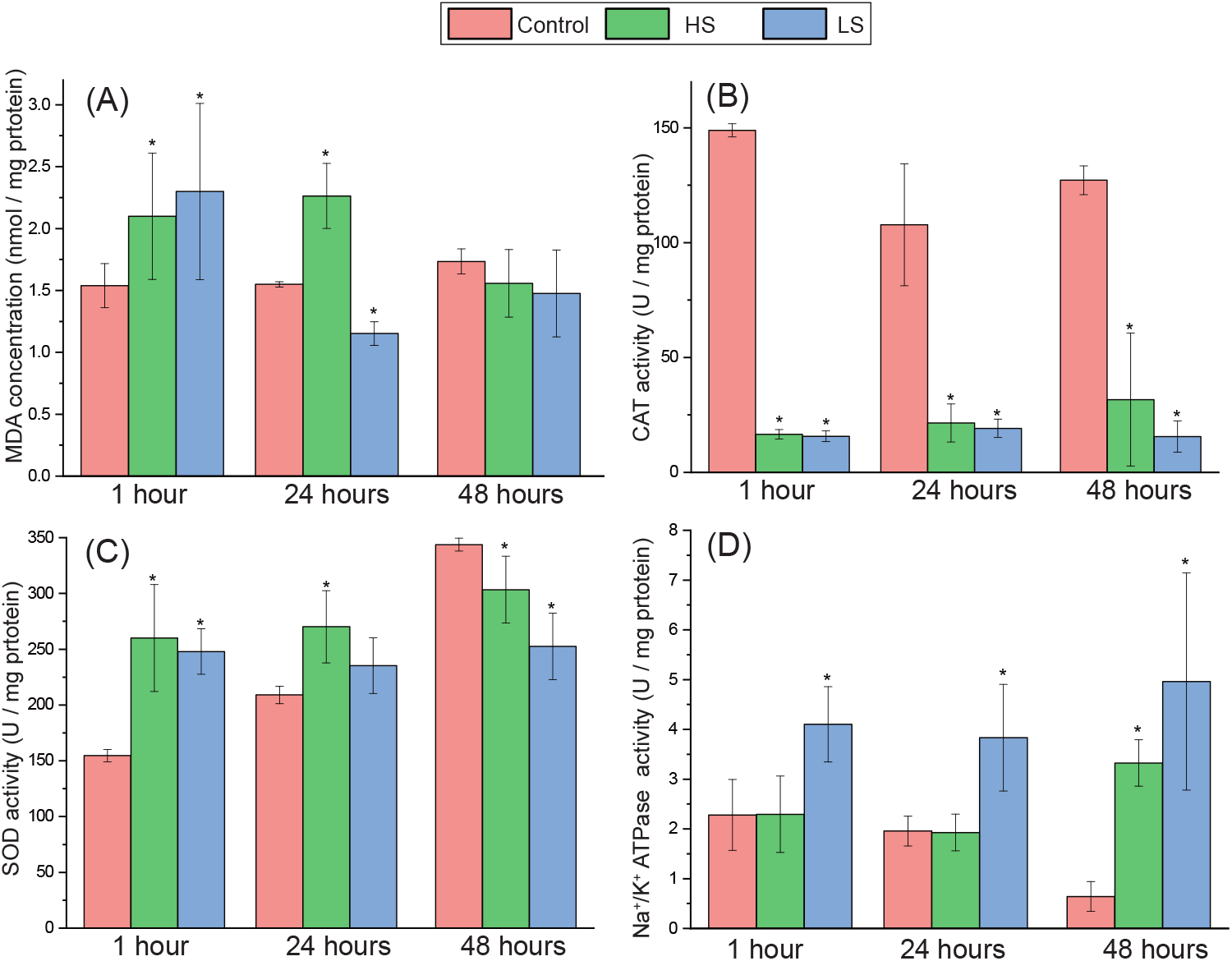
Time course physiological response to high and low salinity stresses in *Ciona savignyi*. Four basic physiological indices include the oxidant stress indicated by MDA (A), antioxidant activity of CAT (B) and SOD (C), and ATP-dependent ion pump activity of Na^+^/K^+^ ATPase (D). The asterisk “*” indicates the statistical difference (*P* < 0.05) between the treated and corresponding control groups.

### Gene expression response to salinity stresses

An average of 19.68 million clean reads with high sequence quality per sample were obtained, and the mapping rate per library ranged from 76% to 87% with an average of 81.72%, resulting in at least 10 million mapped reads per sample for subsequent analyses (Table S1). Based on all 12,172 annotated genes in the reference genome, the overall transcriptomic profile from principal component analysis (PCA) analysis showed that, although variation among individuals could not be explained by salinity changes or durations alone, the low salinity-treated ascidians, particularly for those at 24 and 48 hours, were more separated from the unstressed samples than high salinity-challenged ascidians (Figure S1), indicating LS-induced stronger transcriptomic reprogramming.

After salinity decreased from 30‰ to 20‰, 101, 1172, and 651 genes underwent significant gene expression changes after 1, 24, and 48 hours respectively, while 241, 103, and 264 genes significantly changed their expression when salinity increased from 30‰ to 40‰ (Figure 2A). The Mann-Whitney *U* test (GO-MWU) analysis was performed to explore the functional representation of salinity stress response genes. The immediate response to LS and HS at 1 hour was dominated by up-regulating signal transduction and potassium ion transport functions but repressing biosynthetic process (Figure S2A and S3A). By 24 hours, vesicle-mediated transport process was significantly up-regulated under LS challenge, and notably neurotransmitter transport process showed an opposite direction of gene expression regulation to LS and HS (Figure S2B and S3B). After 48 hours of HS, the initial down-regulation of ribonucleoprotein complex assembly process switched to up-regulation (Figure S2C), but no biological processes were significantly enriched at LS48.

**Figure 2.**
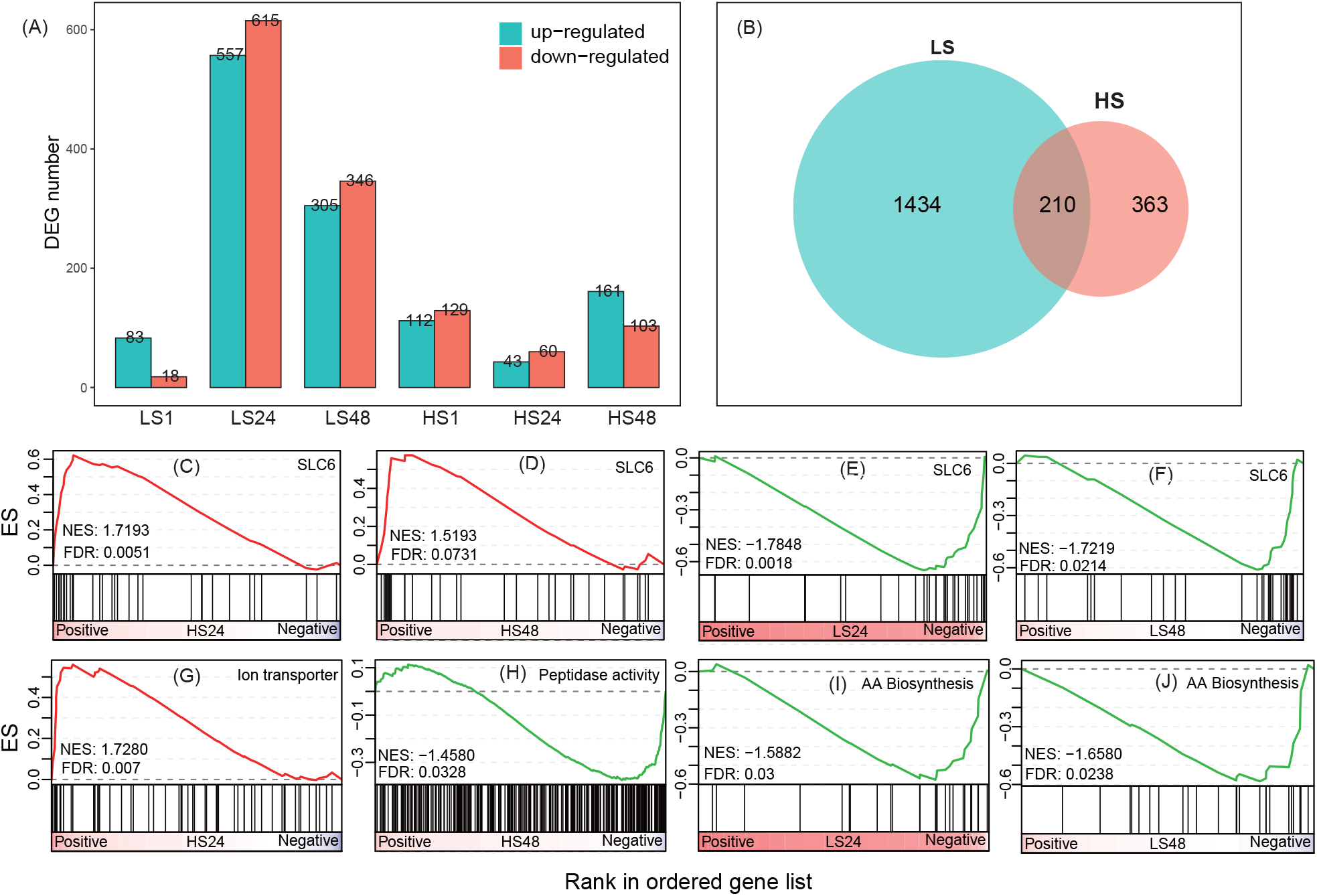
Differential expression induced by high salinity (HS) and low salinity (LS) stresses. (A) The number of up- and down-regulated differentially expressed genes (DEGs) at 1, 24 and 48 hours after two types of salinity treatments; (B) Venn diagram showing common differentially expressed genes (DEGs) and stress-specific DEGs; (C-J) GSEA plots showing significant enrichment of different gene sets (|NES| > 1 and FDR < 0.05), including up-regulation of *SLC6* gene family at HS24 (C), HS48 (D), down-regulation of *SLC6* gene family at LS24 (E) and LS48 (F), up-regulation of ion transporter at HS24 (G), down-regulation of peptidase activity genes at HS48 (H), as well as down-regulation of amino acid (AA) biosynthesis process related genes at LS24 (I) and LS48 (J).

Among 1,644 LS-responsive and 573 HS-responsive differentially expressed genes (DEGs), only 210 DEGs simultaneously responded to HS and LS (Figure 2B). To further clarify the differential response to HS and LS, the gene set enrichment analysis (GSEA) against several osmolyte solutes regulating processes revealed that solute carrier proteins 6 (SLC6) gene family members were significantly enriched among the up-regulated genes at HS24 and HS48 (Figure 2C and 2D), while among the down-regulated genes at LS24 and LS48 (Figure 2E and 2F), indicating free amino acid (FAA) accumulation and FAA loss responding to high salinity and low salinity stresses, respectively. Moreover, ion transmembrane transporting process was significantly stimulated at HS24 while peptidase activity mediated proteolysis was inhibited at HS48 (Figure 2G and 2H). Amino acid biosynthetic process was significantly enriched among down-regulated genes at LS24 and LS48 (Figure 2I and 2J), potentially contributing to the decrease of cellular osmolyte solutes to cope with ambient low salinity.

### Hub genes responding to salinity stresses

To identify key hub/driver genes in transcriptional regulatory networks of salinity stress response, the weighted gene correlation network analysis (WGCNA) was performed using all expressed genes, and each module represents a cluster of genes exhibiting co-expression profiles across all samples. A total of 13 modules were identified with module size ranging from 42 to 2,161 genes. The relationship between module eigengene (the first principal component of gene expression value) and salinity gradients showed that the blue module was the most negatively correlated with salinity gradients, while lightcyan module was positively correlated with salinity gradients (Table S2). The gene expression in blue module was significantly induced by the low salinity stress but suppressed by the high salinity stress (Figure 3A). Hub genes (top 30 genes with high intramodular connectivity) in this module contained transmembrane transport-related genes *SLC49A3* and *mup-4*, cell growth related gene *IGFBP*, methyltransferase gene *METTL27*, two zinc metalloproteinase genes *Nas-13* and *Nas-16*, as well as an adhesion molecule *OTOA* (Figure 3C). All hub genes is listed in detail in Table S3.

**Figure 3.**
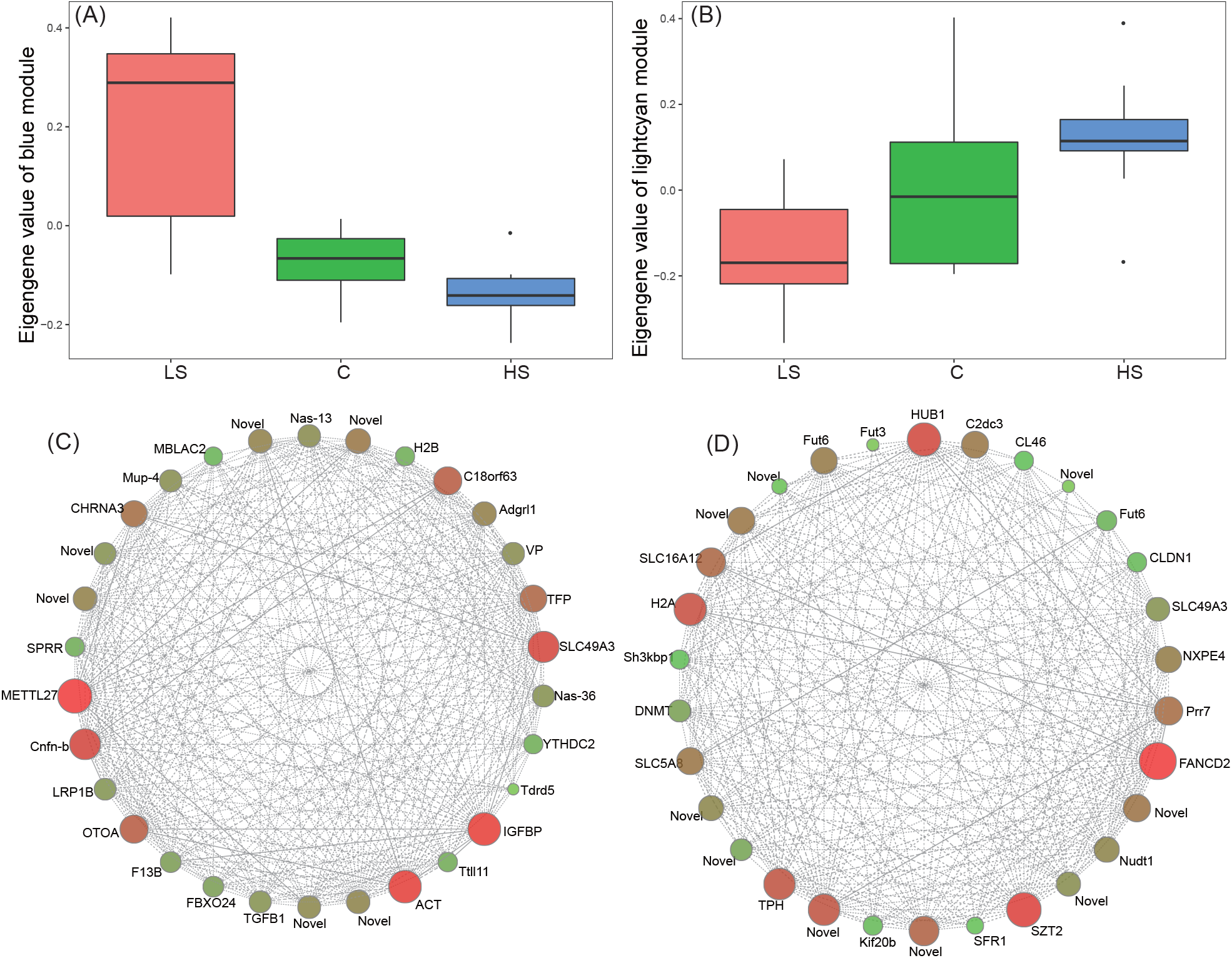
Gene co-expression analysis to identify hub genes responsible for salinity challenges. Eigengene value distribution of blue (A) and lightcyan (B) modules show the gene expression level in high salinity (HS), low salinity (LS), and control groups. Network visualization of gene interactions of top 30 with highest connectivity is shown in blue (C) and lightcyan (D) modules. Each node represents an expressed gene labelled with gene symbol and its detailed information is supplemented in Table S3.

Contrary to the expression profile of blue module, the gene expression in lightcyan module was significantly induced by the high salinity stress but suppressed by the low salinity stress (Figure 3B). The hub genes in lightcyan module included three solute carrier proteins (SLC) family members *SLC16A12, SLC49A3*, and *SLC5A8*, two fucosyltransferase genes *FUT3* and *FUT6*, oxidative stress response gene *FANCD2*, and several protein processing genes such as *HUB1, SZT2* and *C2dc3* (Figure 3D and Table S3). In addition to the annotated genes, many novel genes with unknown functions were also identified as hub genes in these two modules.

### Differentially alternative spliced genes (DASG) and differentially expressed APA (DeAPA) genes in response to salinity stresses

Five types of alternative splicing (AS) events were detected using rMATS, including skipped exon (SE), retained intron (RI), alternative 5’ and 3’ splice site (A5’SS and A3’SS), and mutually exclusive exon (MXE). SE was the most dominant AS type in response to salinity stresses, accounting for 70.66% of all splicing events, followed by MXE (11.75%), A5’SS (10.38%), A3’SS (6.96%), and RI (0.24%) (Figure 4A). Accordingly, the subsequent analyses mainly focused on salinity stress-induced SE changes. After salinity decreases, the inclusion level of 114, 147, and 170 exons changed significantly at 1, 24, and 48 hours, affecting the isoform composition of 44, 101, and 77 genes, respectively, while the high salinity stress significantly induced the inclusion level changes (present spliced in, ΔPSI) of 76, 110, and 81 exons in 46, 48, and 54 genes accordingly (Figure 4B). Among these differentially alternative splicing events (DASEs), the number of obtaining (ΔPSI > 0) and losing (ΔPSI < 0) a particular exon after salinity changes was comparable (Figure S4), and we did not observe a clear tendency of salinity stress induced exon loss or retain. The induced DASGs varied considerably over time, with only seven and three DASGs shared in all three time points after low salinity and high salinity stresses, respectively (Figure 4C), indicating dynamic AS response to salinity challenges. Among the seven DASGs persistently undergone AS regulation in response to low salinity stress, one noteworthy gene was annotated as ABC transmembrane type -1 domain-containing protein (*Abcc1*) which has the ATPase-coupled transmembrane transporter activity and functions in transporting various substrates. While *DDX46* gene was among the three common DASGs in response to the high salinity stress, which is a probable ATP-dependent RNA helicase and plays essential roles in the splicing process.

**Figure 4.**
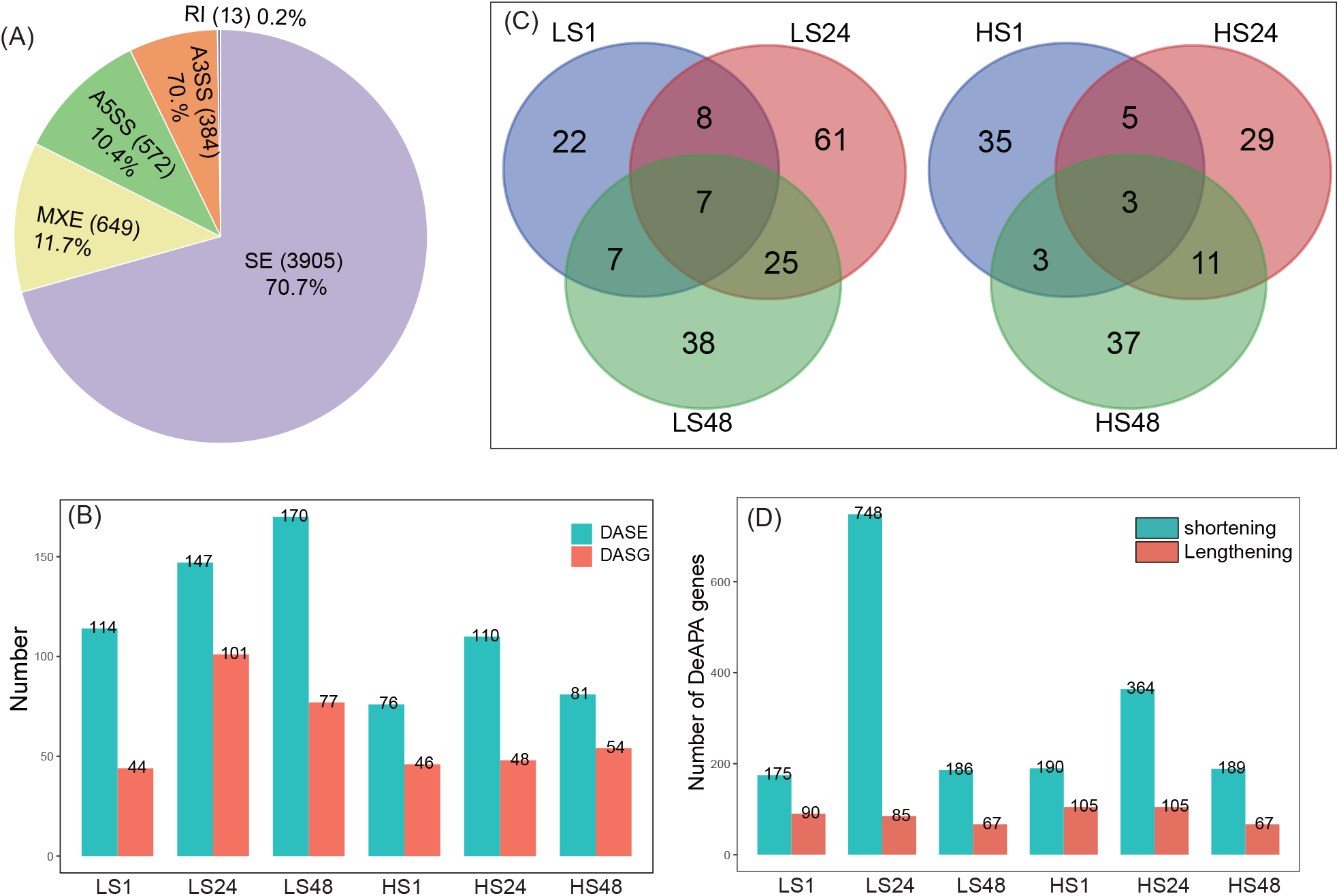
Differentially alternative splicing and alternative polyadenylation profiles in response to salinity stresses. (A) Average percentages of five main types of alternative splicing events between all pairwise comparisons of treatment and control groups detected by rMATS; (B) The number of differentially alternative splicing genes (DASE) and differentially alternative spliced genes (DASG) in each treatment-control comparison group; (C) Venn diagram showing DASGs at different durations under low and high salinity stresses respectively; (D) The number of differentially alternative polyadenylation (DeAPA) genes with shortening and lengthening 3’UTR after salinity stress.

With APAtrap, we identified 7,545 genes with alternative poly (A) sites among all samples, accounting for 62% of all genes in the reference genome. After salinity changes, 1,179 and 808 DeAPA genes significantly changed their poly (A) sites usage in response to low salinity and high salinity stresses, respectively, with 403 genes commonly responding to both stresses. At all three time points of low salinity or high salinity stresses, we detected more salinity stress-induced shortening genes than lengthening genes (Figure 4D), indicating a global 3’UTR shortening profile caused by preferential usage of proximal poly(A) sites. To further test whether salinity stress-induced 3’UTR shortening was related with enhanced gene expression abundance, we compared the gene expression changes of shortening and lengthening gene sets, but we did not detect significantly different gene expression response (Figure S5).

### Stress-induced isoform switch events

Two significant isoform switch events were identified using 3D RNA-seq APP - *SLC2a5* (Gene ID: ENSCSAVG00000011114, Solute carrier family 2, facilitated glucose transporter member 5) and *Cyb5r3* (Gene ID: ENSCSAVG00000011164, NADH-cytochrome b5 reductase 3) (Figure 5). Two alternative transcript isoforms were transcribed from *SLC2a5* gene, which functions as a fructose transporter in salt uptake, isoform 1 (transcript ID: ENSCSAVT00000019133) with 10 exons and isoform 2 (transcript ID: ENSCSAVT00000019134) with nine exons (Figure 5A). Under normal conditions, isoform 2 was the dominant transcript, but its expression was downregulated and the ratio of isoform 1 increased and became the dominant transcript under both high salinity and low salinity stresses (Figure 5B). By comparing the transcript and deduced protein structures, we found that the second exon of isoform 1 was skipped in isoform 2, leading to one more transmembrane region predicted in deduced protein sequence from isoform 1 than that from isoform 2 (Figure 5C). Such change could potentially affect the capacity of fructose transportation and play a role in the osmatic regulation under salinity stresses.

**Figure 5.**
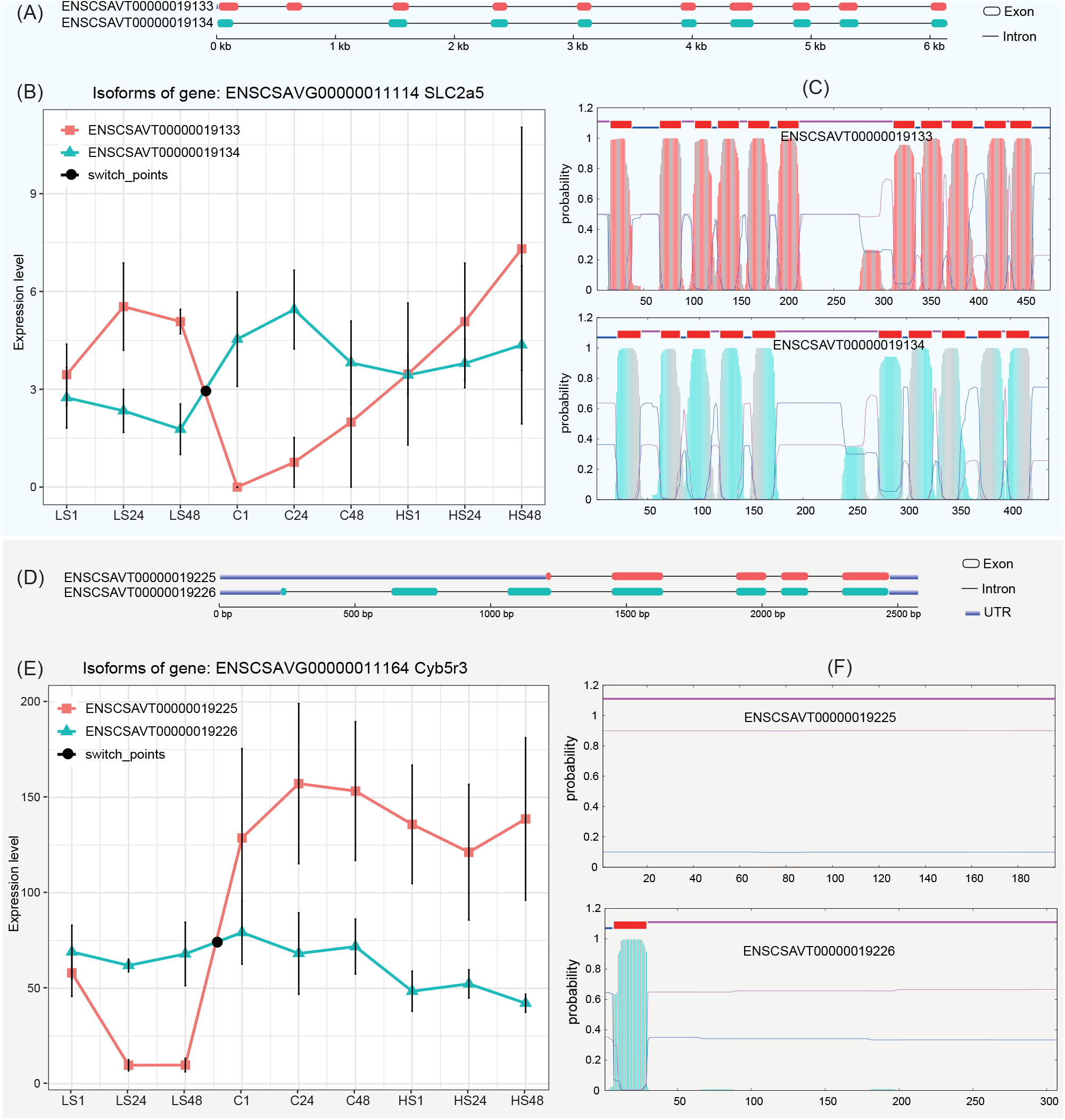
Two isoform switch events occurring in *SLC2a5* and *Cyb5r3* genes. (A) The gene structure of two transcript isoforms of *SLC2a5* gene; (B) The expression changes of two transcript isoforms of *SLC2a5* gene across all the experimental groups; (C) Transmembrane region prediction of two transcript isoforms of *SLC2a5* gene; (D) The gene structure of two transcript isoforms of Cyb5r3 gene; (E) The expression changes of two transcript isoforms of Cyb5r3 gene across all the experimental groups; (F) Transmembrane region prediction of two transcript isoforms of Cyb5r3 gene.

Another significant isoform switch event occurred in *Cyb5r3* gene, leading to two alternative transcripts with different 5’ coding regions (Figure 5D): isoform 1 (transcript ID: ENSCSAVT00000019225) and isoform2 (transcript ID: ENSCSAVT00000019226). Under control condition and high salinity stress, isoform 1 was the dominant transcript, while it was downregulated under low salinity stress, making isoform 2 as the dominant one (Figure 5E). By predicting the transmembrane region of deduced proteins, the appearance of transmembrane region in isoform 2 made it a transmembrane protein locating at endoplasmic reticulum membrane or mitochondrion outer membrane (Figure 5F), while the lack of transmembrane region in isoform 1 made it a soluble protein which mainly functions in cytoplasm.

### Complementary regulations of AS, APA, and gene expression mechanisms

Overall, the number of genes undergoing AS regulation was far fewer than the genes with mRNA abundance changes in response to salinity shifts, with all splicing abundance ratio (SAR) lower than 100 (Figure 6A). While the number of genes undergoing APA regulation far exceeded that of DEGs at LT1 and HS24 (polyadenylation abundance ratio PAR > 100, Figure 6A), suggesting the dominant role of APA mechanism in transcriptomic plasticity at those time points. By comparing the genes involved in gene expression, AS and APA regulations in response to salinity stresses, we found relatively low proportion of shared genes coordinately acted on more than two regulatory layers, with only four and zero genes shared by all three regulations under low salinity and high salinity stresses, respectively (Figure 6B and 6C). Most of the DASGs and DeAPA genes were not differentially expressed, and most of DeAPA genes were not differentially alternatively spliced, indicating relatively independent regulations among AS, APA, and gene expression plasticity. To test whether gene structure affected the choice of different regulatory mechanisms, we found that the genes with more exons tended to adopt AS changes to respond to salinity challenges (Figure 6D and 6E). Different regulatory mechanisms acted on varied genes and corresponding biological functions (Figure S6), and all these enriched functions were important for *C. savignyi* acclimation to salinity changes. For example, ion transmembrane transport related genes changed gene expression to cope with decreased salinity, and genes responding to stimulus were subjected to AS regulation, while organic substance transport related genes adopted APA regulation (Figure S6). Although the three response mechanisms affected different genes, there exist some interactions among them, e.g., the genes associated with mRNA poly(A) tail shortening were alternatively spliced, and RNA splicing relevant genes were under both gene expression and APA regulations in response to low salinity stress (Figure S6A).

**Figure 6.**
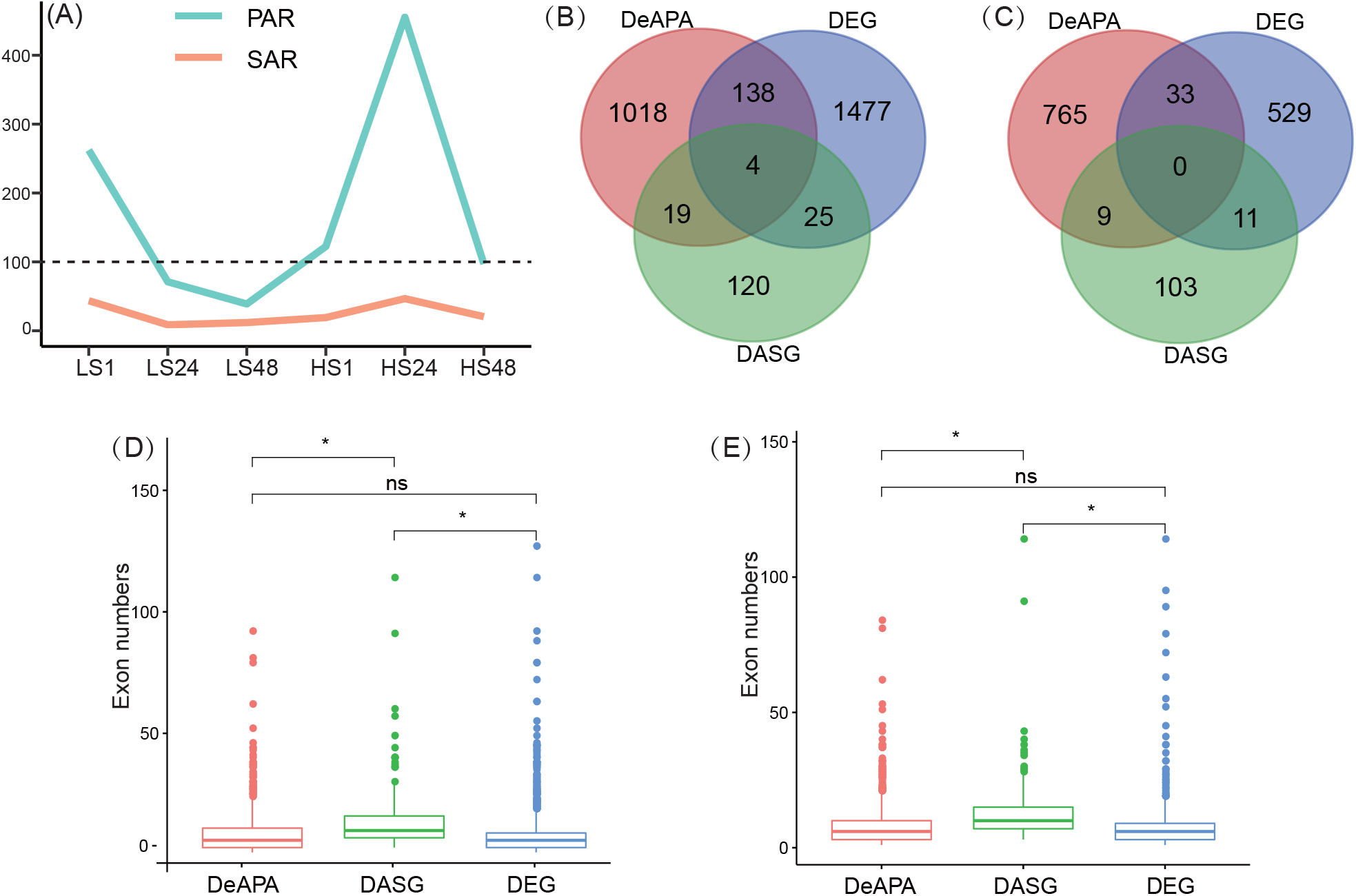
Comparisons of genes underlying three different regulatory mechanisms including gene expression, alternative splicing, and alternative polyadenylation. (A) SAR (splicing abundance ratio) and PAR (polyadenylation abundance ratio) in different treatment groups, indicating the number of alternative splicing changes or poly(A) changes relative to DEGs; (B) Venn diagram showing DEG, DASG and DeAPA under LS and HS stress (C); (D) Exon number distributions of the genes regulated by three different mechanisms under LS and HS stress (D) respectively, the asterisk “*” indicates the statistical difference (*P* < 0.05), “ns” indicates non-significant.

## DISCUSSION

Exploring the complex mechanisms underpinning environment-induced plastic response is an important and prerequisite step for understanding organismal performance in rapidly changing environments. Post-transcriptional processes such as AS and APA regulations have been widely overlooked in organismal response to environmental changes (Salisbury et al. 2021; Verta and Jacobs 2022). In this study, we used a model marine invasive species *C. savignyi* to investigate multiple layers of plastic response to salinity shifts, which simulated the real salinity changes during several days’ transoceanic voyages of the invasion process (Klein et al. 2010; Bereza and Shenkar 2022). Our results showed that different layers of plastic response were rapidly and dynamically induced to cope with salinity changes. Based on the number of affected genes, initial plasticity was dominated by gene expression plasticity at most time points under high salinity and low salinity stresses, except for APA plasticity as dominant mechanism at LS1 and HS24. We also found that gene expression, AS, and APA plastic response acted on different genes and biological functions, playing complementary roles in rapid salinity acclimation. Interestingly, the choice of target genes by different mechanisms was related with gene structures, for example, genes under AS regulation tended to have more exons. Moreover, we observed partly different plastic response induced by low salinity and high salinity stresses, and identified key biological functions and candidate genes involved in salinity acclimation. Altogether, the results obtained here reveal the complex interplays among different initial plastic response to the time course of salinity challenges, providing novel insights into the roles of complex molecular plasticity towards adaptive evolution.

### Interplay of multidimensional plasticity under salinity stresses

Interestingly, gene expression, AS, and APA plastic response affected different genes and biological functions, indicating their complementary regulatory roles in rapid salinity acclimation. The relatively independent roles of gene expression and AS were also reported in the parallel adaptive evolution of salmonid fish (Jacobs and Elmer 2021), environmentally determined phenotypes of pea aphid (Grantham and Brisson 2018), acute copper stress response of *Daphnia* (Suresh et al. 2020), and short-term maintaining mineral nutrient homeostasis of rice (Dong et al. 2018). Consistently with our results, there were more genes undergoing gene expression changes than AS changes in most studies (Grantham and Brisson 2018; Shalgi et al. 2014; Suresh et al. 2020; Tian et al. 2020; Huang and Zhan 2021). However, a few studies demonstrated contrasting results with more or comparable genes undergoing AS regulation (Singh et al. 2017; Dong et al. 2018; Jacobs and Elmer. 2021), indicating that their relative importance varied among species, types of environmental stresses, and challenge durations. Compared with AS, studies on APA plastic response to environmental changes and its relationship with gene expression or AS have just emerged in recent years. Several studies demonstrated that 3’UTR landscape was reshaped under environmental stresses through the APA mechanism (Zheng et al. 2018; Sadek et al. 2019; Ye et al. 2019). Consistently with findings here, those studies detected dominant shortening profiles of 3’UTRs. Such environmentally induced 3’UTR shortening can lead to a higher gene expression level because shortened 3’UTR might lack some miRNA target sites and further escape from mRNA degradation processes (Ye et al. 2019; Tian and Manley 2017). However, we did not detect such relationship between 3’UTR shortening and higher gene expression, and the low overlap between DEGs and DeAPA genes also indicated independent roles of these two mechanisms. Moreover, the number of DeAPA genes far exceeded DEGs at LS1 and HS24, suggesting important roles of APA mechanism in specific phases of salinity acclimation. In addition, even though compelling evidence showed AS and APA as coordinated mechanisms (Nazim et al. 2018), our results, particularly the extremely low overlap between the affected genes, support independent roles of AS and APA during the plastic response to salinity challenges.

One possible explanation for independent relationship of gene expression, AS, and APA mechanisms is that they are controlled by different regulatory machineries involving distinct cis-elements and trans-acting factors, such as promoters and transcription factors for gene expression, splice sites, and spliceosome for AS, and polyadenylation sites and cleavage factors for APA (Schaefke et al. 2018; Sadek et al. 2019). Meanwhile, we also found a certain extent of interactions among different regulatory layers, for example, RNA splicing machinery related genes were differentially expressed, and APA machinery related genes were alternatively spliced under the low salinity stress. Despite of their subtle relationship, the affected biological functions such as transmembrane transporting, stress response, biosynthesis or metabolic processes of amino acid and fatty acid have been widely reported in rapid salinity acclimation in other species (Posavi et al. 2020; Niu et al. 2020; Maynard et al. 2018; Jeffries et al. 2019). Therefore, AS and APA-based plasticity response should provide additional layers of initial plasticity to gene expression, jointly contributing to rapid response to salinity stresses.

### Contrast between plastic response to high salinity and stresses

Salinity fluctuates dramatically during trans-oceanic voyages (Ghabooli et al. 2016; Bereza and Shenkar 2022), posing environmental constraints for the survival and establishment of invasive species. As osmoconformers, ascidians must be able to adjust their intercellular osmolarity when ambient salinity changes (Sokolov and Sokolova 2019). Osmoregulatory mechanisms of osmoconformers have been extensively assessed in mollusks, and similarly to results obtained in *Ciona* here, free amino acids (FAAs) were the preferential osmolytes to regulate osmolarity (Liu et al. 2018; Pourmozaffar et al. 2020; Sokolov and Sokolova 2019; Yancey, 2005). The *SLC6* gene family is responsible for transporting amino acids, nutrients, or other kinds of osmolytes (Bröer and Gether 2012), the expansion of which was found in the genome of *C. savignyi* (Ren et al. 2019). Our results showed that *SLC6* genes were significantly up-regulated under the high salinity stress and down-regulated under the low salinity stress, as well as the down-regulation of amino acid biosynthetic process under the low salinity stress, consistently supporting that FAA, as the main osmolytes, accumulated to increase intracellular osmolarity to cope with the ambient high salinity stress while decreased under the low salinity stress. Besides, some studies argued that inorganic ions also participated in salinity acclimation of osmoconformers, but in fact the relative contributions of inorganic ions and FAA largely varied among species, tissue types, stress intensity, and duration time. For examples, the Pacific oyster *Crassostrea gigas* transcriptionally triggered the FAA biosynthesis process to cope with salinity stresses (Zhao et al. 2016), while the white clam *Amarilladsma mactroides* mainly recruited Na^+^ and very low amounts of FAAs and the lagoon cockle *Erodona mactroides* only used Na^+^ and K^+^ as effectors for osmoregulation (Medeiros et al. 2020). We observed in this study that ion transmembrane transporting related genes were significantly up-regulated at HS48, indicating that their involvement in high salinity response might be environment-specific or time-dependent. However, more direct evidence such as the time-course changes of cellular osmolarity, ion (e.g., Na^+^ and K^+^) or FAA (e.g., taurine, glycine, alanine, and proline, etc.) concentration under salinity stresses should be provided to illustrate the dynamic osmoregulatory process in the further studies.

### Key candidate genes involved in response to salinity stresses

SLC genes are a group of over 400 membrane proteins to transport extraordinarily diverse of solutes including organic molecules, inorganic ions, and gas ammonia (He et al. 2009). Different SLC genes were found to be involved in response to salinity stresses, such as *SLC39A6* and *SLC5A9* in the tiger puffer *Takifugu rubripes* (Jiang et al. 2020), *SLC6A6* in *Cynoglossus semilaevis* (Si et al. 2018), and *SLC4A4* and *SLC12A2* in hybrid tilapia (Su et al. 2020). Our present study found that SLC gene family members were consistently identified as hub genes on gene expression regulatory layer using WGCNA analysis, including *SLC49A3, SLC5A8*, and *SLC16A12*, indicating their important roles in salinity acclimation of ascidians. Moreover, we also identified a key isoform switch event occurred on another SLC gene *SLC2a5. SLC2a5* gene is a member of GLUT transporter specific to fructose (Burant et al. 1992), and its expression change has been linked to several metabolic disorders and human cancers (Barone et al. 2009). A total of 12 transmembrane regions (TMs) are necessary for its normal function, and incomplete TMs might impair its transporting activities (Mueckler and Makepease 2002). In the present study, we found that the shorter isoform of *SLC2a5* with 11 TMs was constitutively expressed in the control group, while the longer isoform with 12 TMs was induced under both salinity stresses, indicating the facilitated fructose transporting function for osmoregulation under salinity stresses.

In conclusion, our results revealed that rapid plastic response patterns varied greatly with salinity shift context (HS and LS), short timescales (1, 24, and 48 hours), and distinct regulatory mechanisms (gene expression, AS, and APA), and such multidimensional plastic response involves different genes and biological functions. Considering that different layers of molecular plastic response will be jointly translated to functional protein changes and further phenotypic diversity that natural selection can act upon, how to integrate all different plastic response or discern the dominant plastic response, can offer valuable insights into the predictability of organismal adaptation to environmental changes.

## MATERIALS AND METHODS

### Organism collection and experimental design

*Ciona savignyi* individuals were sampled from the aquaculture scallop cages on the coast of Dalian, Liaoning Province, China (38°49’13’’N, 121°24’20’’E). All collected ascidians were immediately transported to the laboratory and then acclimated in aerated tanks with ambient temperature of 15 ± 1 °C and salinity of 30‰ (the measured field parameters) for one week. Following acclimation, individuals were randomly assigned to one of the three following groups: control (C), high salinity (HS, 40‰), and low salinity (LS, 20‰), with all other environmental factors unchanged. The target salinities were achieved by adding instant sea salt or dechlorinated tap water. Considering that pharynx covered with gill slits is the primary interface where ambient water current and metabolic waste directly enter and exit, the pharynx tissues of six replicate ascidians were sampled individually after 1, 24, and 48 hours of stress exposure according to our previous study (Huang et al. 2017). All samples were immediately preserved in liquid nitrogen and then stored at -80 °C.

### Assessment of physiological indicators

We chose four basic physiological indices, including MDA indicating lipid peroxidation of membranes, the enzyme activity of SOD and CAT indicating antioxidant status, and the activity of Na^+^/K^+^ ATPase indicating ion transport regulation, to study the dynamic physiological response to salinity stresses, with six ascidian individuals as biological replicates in each treatment group. The four indices were measured and calculated according to the kit instructions (Jiancheng, Nanjing, China).

### RNA extraction, cDNA library construction, and RNA sequencing

Total RNA was extracted from 50-100 mg pharynx tissue of each sample using Trizol reagent (Ambion, USA) and then treated with DNase (Promega, USA) to remove potential DNA contamination. Three best RNA samples were selected based on RNA integrity (RNA integrity number ≥ 7) and purity, which were evaluated using 1.5% (w/v) agarose gel electrophoresis and Agilent 2100 Bioanalyzer (Agilent Technologies, USA). A total of 27 ascidian samples (three treatments × three time points × three biological replicates) were used for cDNA library construction using the NEBNext^®^ UltraTM RNA Library Prep Kit (New England Biolabs, USA) and RNA sequencing on the Illunima Hiseq 4000 sequencing platform with the pair end of 150 bp strategy. All raw sequencing data was deposited in the National Center for Biotechnology Information (NCBI) Sequence Read Archive (SRA) database under the accession number SRP152910.

### Data processing and DEGs identification

Raw reads were firstly processed to filter out the adaptor and low-quality sequence using Trimmomatic version 0.36 (Bolger et al. 2014), and the acquired clean reads were then mapped to the *C. savignyi* reference genome CSAV 2.0 using HISAT2 with the following parameters: -p 8 -N 1 -L 20 -i S, 1, 0.5 -D 25 -R 5 –mp 1, 0 -sp 3, 0 -x hisat2_index (Kim et al. 2015). The mapped reads were subsequently assembled to the transcripts and the abundance of gene expression was estimated using StringTie software with default settings (Pertea et al. 2015). The read count number of each gene was normalized using the relative log (rlog) transformation method. Genes with read count lower than 10 in more than 90% samples were excluded from further analyses. DEGs were identified by pairwise comparing gene expression between each environmental treatment group and its corresponding control group at the same time point using the R package DESeq2 (Love et al. 2014), including LS1 *vs*. C1, LS24 *vs*. C24, LS48 *vs*. C48, HS1 *vs*. C1, HS24 *vs*. C24, HS48 *vs*. C48. The genes with the fold change of expression level larger than two as well as the false discovery rate (FDR) lower than 0.05 were considered as potential DEGs. The visualization of variation among all samples based on rlog transformation normalized read counts of all expressed genes was conducted by principal component analysis (PCA).

### Weighted gene co-expression network analysis

To explore the potential regulatory relationships among all genes in response to salinity stresses, WGCNA was constructed using the WGCNA package in R (Langfelder and Horvath 2008). According to the tutorials for WGCNA package, the optimal soft threshold (power = 16) was used for calculating an adjacency matrix based on the scale-free topology criterion using the pickSoftThreshold function of WGCNA. Co-expression modules were then constructed by one-step network construction method. The Spearman correlation between module eigengene and salinity gradients was calculated, and significantly correlated modules (the absolute value of correlation coefficient > 0.5, *P* value < 0.01) were considered as salinity stress response modules. The top 30 hub genes in the given module were selected by intramodular connectivity, and the interactions among those genes in salinity responsible modules were visualized by OmicShare tools (http://www.omicshare.com/tools).

### Alternative splicing analysis

rMATS v.4.0.2 software was used to detect DASE of five different AS types, including SE, RI, A5’SS and A3’SS, and MXE. rMATS quantified the inclusion level of certain alternative splicing site (present spliced in, PSI or φ) between treatment and corresponding control groups (ΔPSI or Δφ) (Shen et al. 2014). DASEs were defined as those with |ΔPSI| > 10% and adjusted *P* value < 0.05. Splicing abundance ratio (SAR) was calculated by DASG (differentially alternative splicing gene) / DEG number × 100, indicating the extent to which AS regulation dominate the transcriptome changes during acclimation to salinity stresses (Habowski et al. 2020). Considering that rMATS detected DASEs at the exon level, the significant isoform switch (IS) events among treatment groups were detected using the 3D RNA-seq APP at the transcript isoform level (Guo et al. 2020). The gene structures of identified IS genes were illustrated using Gene Structure Display Server 2.0 (GSDS, http://gsds.cbi.pku.edu.cn/) by aligning the coding sequences with the corresponding genomic DNA sequences from the same genes. The potential transmembrane regions in proteins deduced from different transcript isoforms were predicted by TMHMM server v2.0 tool (http://www.cbs.dtu.dk/services/TMHMM/).

### Alternative polyadenylation (APA) analysis

APAtrap was used to detect potential APA sites and differential APA events between treatment and corresponding control groups at the same time point (Ye et al. 2018). Differentially expressed APA (DeAPA) genes were defined according to the following three parameters: percentage difference (PD, the difference of APA sites usage between two samples) greater than 0.20, FDR lower than 0.05, as well as the positive or negative of r value (Pearson product-moment correlation coefficient, in the present study the positive *r* value indicates shorter 3’UTR whereas the negative *r* value indicates longer 3’UTR of the treatment group compared with the corresponding control group). Analogous to SAR, polyadenylation abundance ratio (PAR) was calculated by DeAPA / DEG number × 100, indicating the extent to which APA regulation dominate the gene expression changes (Habowski et al. 2020).

### Gene functional enrichment analysis

Three different analyses were conducted to explore the biological functions of salinity responsive genes. For the genes changing their expression level to cope with salinity shifts, we used Mann-Whitney *U* test (GO-MWU) to screen the significantly up- or down-regulated GO categories (https://github.com/z0on/GO_MWU), based on the rank of log *P* values from above DEseq2 analysis (Wright et al. 2015). In order to assess the roles of several key biological processes regulating cellular osmolyte solutes for osmoconformers (Pourmozaffar et al. 2020), such as solute carrier proteins 6 family (*SLC6*) related to FAA transport, ion transmembrane transport, amino acid biosynthesis, proteolysis which were selected according to GO annotation (Table S4), GSEA was performed on gene expression data with GSEA v.4.1.0 program (Subramanian et al. 2005), using the pre-ranked method with log2 fold change value as gene ranking statistic. Gene lists with the absolute value of normalized enrichment score (NES) > 1.0 and FDR < 0.05 were considered significant. For the functional comparison among DEGs, DASGs and deAPA genes, Gene ontology (GO) functional enrichment analysis was conducted by Hypergeometric test using online Omicshare CloudTools (http://www.omicshare.com/tools). Removing redundancy of the significantly enriched GO terms was then conducted by online tool REVIGO (http://revigo.irb.hr).

## Data availability

All raw sequencing data was deposited in the National Center for Biotechnology Information (NCBI) Sequence Read Archive (SRA) database under the accession number SRP152910.

## Author Contributions

X.H., N.S. and A.Z. conceived this study and designed the experiment. X.H. and H.L. conducted the experiments and analyzed the data. X.H. wrote the manuscript. All authors reviewed and commented on the manuscript.

## FIGURE LEGENDS

**Figure S1**. Principal Component Analyses (PCA) of all expressed genes. The numbers in parentheses indicate the proportion of variance explained by that PCA dimension.

**Figure S2**. GO categories significantly enriched with up- and down-regulated Differentially Expressed Genes (DEGs) using Mann-Whitney *U* test under HS stress.

**Figure S3**. GO categories significantly enriched with up- and down-regulated Differentially Expressed Genes (DEGs) using Mann-Whitney *U* test under LS stress.

**Figure S4**. The ΔPSI distributions of DASE in different treatment groups under salinity stresses.

**Figure S5**. Log2 Foldchange distributions of DeAPA genes in different treatment groups.

**Figure S6**. GO enrichment analysis for three subsets of genes under HS (A) and LS (B) stress respectively.

## REFFERENCES

Barone S, Fussell SL, Singh AK, Lucas F, Xu J, Kim C, Wu X, Yu Y, Amlal H, Seidler U, et al. 2009. Slc2a5 (Glut5) is essential for the absorption of fructose in the intestine and generation of fructose-induced hypertension. J Biol Chem 284(8): 5056–5066. doi:10.1074/jbc.M808128200

Bereza D, Shenkar D. 2022. Shipping voyage simulation reveals abiotic barriers to marine bioinvasions. Sci Total Environ 837: 155741. doi:10.1016/j.scitotenv.2022.155741

Bock DG, Kantar MB, Caseys C, Matthey-Doret R, Rieseberg LH. 2018. Evolution of invasiveness by genetic accommodation. Nat Ecol Evol 2(6): 991–999. doi:10.1038/s41559-018-0553-z

Bolger AM, Lohse M, Usadel B. 2014. Trimmomatic: Aflexible trimmer for Illumina sequence data. Bioinformatics 30(15): 2114–2120. doi:10.1093/bioinformatics/btu170

Briski E, Bailey SA, Casas-Monroy O, DiBacco C, Kaczmarska I, Lawrence JE, Leichsenring J, Levings C, MacGillivary ML, McKindsey CW, et al. 2013. Taxon- and vector-specific variation in species richness and abundance during the transport stage of biological invasions. Limnol Oceanogr 58(4): 1361–1372. doi:10.4319/lo.2013.58.4.1361

Bröer S, Gether U. 2012. The solute carrier 6 family of transporters. Brit J Pharmacol 167: 256–278. doi:10.1111/j.1476-5381.2012.01975.x

Burant CF, Takeda J, Brot-Laroche E, Bell GI, Davidson NO. 1992. Fructose transporter in human spermatozoa and small intestine is GLUT5. J Biol Chem 267: 14523–14526.

Chen Z, Huang X, Fu R, Zhan A. 2022. Neighbours matter: Effects of genomic organization on gene expression plasticity in response to environmental stresses during biological invasions. Comp Biochem Phys D 42: 100992. doi: 10.1016/j.cbd.2022.100992

Dong C, He F, Berkowitz O, Liu J, Cao P, Tang M, Shi H, Wang W, Li Q, Shen Z, et al. 2018. Alternative splicing plays a critical role in maintaining mineral nutrient homeostasis in rice (Oryza sativa). Plant Cell 30(10): 2267–2285. doi:10.1105/tpc.18.00051

Fofonoff PW, Ruiz GM, Steves B, Simkanin C, Carlton JT. 2018. National exotic marine and estuarine species information system. http://invasions.si.edu/nemesis/.

Fox RJ, Donelson JM, Schunter C, Ravasi T, Gaitán-Espitia JD. 2019. Beyond buying time: the role of plasticity in phenotypic adaptation to rapid environmental change. Philos T R Soc B 374(1768): 20180174. doi:10.1098/rstb.2018.0174

Fu R, Huang X, Chen Y, Chen Z, Zhan A. 2021. Interactive regulations of dynamic methylation and transcriptional responses to recurring environmental stresses during biological invasions. Front Mar Sci 8: 800745. doi: 10.3389/fmars.2021.800745

Ghabooli S, Zhan A, Paolucci E, Hernandez MR, Briski E, Cristescu ME, MacIsaac HJ. 2016. Population attenuation in zooplankton communities during transoceanic transfer in ballast water. Ecol Evol 6(17): 6170–6177. doi:10.1002/ece3.2349

Grantham M., Brisson JA. 2018. Extensive Differential Splicing Underlies Phenotypically Plastic Aphid Morphs. Mol Biol Evol 35(8): 1934–1946. doi:10.1093/molbev/msy095

Guo W, Tzioutziou N, Stephen G, Milne I, Calixto C, Waugh R, Brown JWS, Zhang R. 2020. 3D RNA-seq - a powerful and flexible tool for rapid and accurate differential expression and alternative splicing analysis of RNA-seq data for biologists. RNA Biol 18: 1574–1587. doi: 10.1080/15476286.2020.1858253.

Habowski AN, Flesher JL, Bates JM, Tsai CF, Martin K, Zhao R, Ganesan AK, Edwards RA, Shi T, Wiley HS, et al. 2020. Transcriptomic and proteomic signatures of stemness and differentiation in the colon crypt. Commun Biol 3(1): 453. doi:10.1038/s42003-020-01181-z

He L, Vasiliou K, Nebert DW. 2009. Analysis and update of the human solute carrier (SLC) gene superfamily. Hum Genomics 3(2): 195. doi:10.1186/1479-7364-3-2-195

Huang X, Li S, Ni P, Gao Y, Jiang B, Zhou Z, Zhan A. 2017. Rapid response to changing environments during biological invasions: DNA methylation perspectives. Mol Ecol 26(23): 6621–6633. doi:10.1111/mec.14382

Huang X, Zhan A. 2021. Highly dynamic transcriptional reprogramming and shorter isoform shifts under acute stresses during biological invasions. RNA Biology 18(3): 340–353. doi:10.1080/15476286.2020.1805904

Jacobs A, Elmer KR. 2021. Alternative splicing and gene expression play contrasting roles in the parallel phenotypic evolution of a salmonid fish. Mol Ecol 30: 4955–4969. doi:10.1111/mec.15817

Jeffries KM, Connon RE, Verhille CE, Dabruzzi TF, Britton MT, Durbin-Johnson BP, Fangue NA. 2019. Divergent transcriptomic signatures in response to salinity exposure in two populations of an estuarine fish. Evol Appl 12(6): 1212–1226. doi:10.1111/eva.12799

Jiang JL, Xu J, Ye L, Sun ML, Jiang ZQ, Mao MG. 2020. Identification of differentially expressed genes in gills of tiger puffer (Takifugu rubripes) in response to low-salinity stress. Comp Biochem Physiol B Biochem Mol Biol 243-244: 110437. doi:10.1016/j.cbpb.2020.110437

Josephs EB. 2021. Gene expression links genotype and phenotype during rapid adaptation. Mol Ecol 11(1): 17–28. doi:10.1111/eva.12528

Kim D, Langmead B, Salzberg SL. 2015. HISAT: a fast spliced aligner with low memory requirements. Nat Methods 12(4): 357–360. doi:10.1038/nmeth.3317

Klein G, MacIntosh K, Kaczmarska I, Ehrman JM. 2009. Diatom survivorship in ballast water during trans-Pacific crossings. Biol Invasions 12(5): 1031–1044. doi:10.1007/s10530-009-9520-6

Komoroske LM, Jeffries KM, Whitehead A, Roach JL, Britton M, Connon RE, Verhille C, Brander SM, Fangue NA. 2021. Transcriptional flexibility during thermal challenge corresponds with expanded thermal tolerance in an invasive compared to native fish. Evol Appl 14: 931–949.doi:10.1111/eva.13172

Lambert CC, Lambert G. 2003. Persistence and differential distribution of nonindigenous ascidians in harbors of the Southern California Bight. Mar Ecol Prog Ser 259: 145–161.

Langfelder P, Horvath S. 2008. WGCNA: an R package for weighted correlation network analysis. BMC Bioinformatics 9: 559. doi:10.1186/1471-2105-9-559

Li H, Huang X, Zhan A. 2020. Stress memory of recurrent environmental challenges in marine invasive species: Ciona robusta as a case study. Front Physiol 11: 94. doi: 10.3389/fphys.2020.00094

Liu Z, Zhou Z, Wang L, Li M, Wang W, Yi Q, Huang S, Song L. 2018. Dopamine and serotonin modulate free amino acids production and Na(+)/K(+) pump activity in Chinese mitten crab Eriocheir sinensis Under Acute Salinity Stress. Front Physiol 9: 1080. doi:10.3389/fphys.2018.01080

Love MI, Huber W, Anders S. 2014. Moderated estimation of fold change and dispersion for RNA-seq data with DESeq2. Genome Biol 15(12): 550. doi:10.1186/s13059-014-0550-8

Maynard A, Bible JM, Pespeni MH, Sanford E, Evans TG. 2018. Transcriptomic responses to extreme low salinity among locally adapted populations of Olympia oyster (Ostrea lurida). Mol Ecol 27(21): 4225–4240. doi:10.1111/mec.14863

Medeiros IPM, Faria SC, Souza MM. 2020. Osmoionic homeostasis in bivalve mollusks from different osmotic niches: Physiological patterns and evolutionary perspectives. Comp Biochem Physiol A Mol Integr Physiol 240: 110582. doi:10.1016/j.cbpa.2019.110582

Metzger DCH, Schulte PM. 2018. Similarities in temperature-dependent gene expression plasticity across timescales in threespine stickleback (Gasterosteus aculeatus). Mol Ecol 27(10): 2381–2396. doi:10.1111/mec.14591

Mueckler M, Makepeace C. 2002. Analysis of transmembrane segment 10 of the Glut1 glucose transporter by cysteine-scanning mutagenesis and substituted cysteine accessibility. J Biol Chemi 277(5): 3498–3503. doi:10.1074/jbc.M109157200

Nazim M, Nasrin F, Rahman MA. 2018. Coordinated Regulation of Alternative Splicing and Alternative Polyadenylation. Journal of Genetics and Genetic Engineering 2(3): 26–34.

Niu J, Hu XL, Ip JCH, Ma KY, Tang Y, Wang Y, Qin J, Qiu JW, Chan TF, Chu KH. 2020. Multi-omic approach provides insights into osmoregulation and osmoconformation of the crab Scylla paramamosain. Scic Rep 10(1): 21771. doi:10.1038/s41598-020-78351-w

Pereira-Castro I, Moreira A. 2021. On the function and relevance of alternative 3’-UTRs in gene expression regulation. Wires RNA e1653. doi:10.1002/wrna.1653

Pertea M, Pertea GM, Antonescu CM, Chang TC, Mendell JT, Salzberg SL. 2015. StringTie enables improved reconstruction of a transcriptome from RNA-seq reads. Nat Biotechnol 33(3): 290–295. doi:10.1038/nbt.3122

Posavi M, Gulisija D, Munro JB, Silva JC, Lee CE. 2020. Rapid evolution of genome-wide gene expression and plasticity during saline to freshwater invasions by the copepod Eurytemora affinis species complex. Mol Ecol 29(24): 4835–4856. doi:10.1111/mec.15681

Pourmozaffar S, Tamadoni Jahromi S, Rameshi H, Sadeghi A, Bagheri T, Behzadi S, Gozari M, Zahedi MR, Abrari Lazarjani S. 2020. The role of salinity in physiological responses of bivalves. Rev Aquacult 12(3): 1548–1566. doi:10.1111/raq.12397

Price J, Harrison MC, Hammond RL, Adams S, Gutierrez-Marcos JF, Mallon EB. 2018. Alternative splicing associated with phenotypic plasticity in the bumble bee Bombus terrestris. Mol Ecol 27(4): 1036–1043. doi: 10.1111/mec.14495

Ren P, Wei J, Yu H, Dong B. 2019. Identification and functional characterization of solute carrier family 6 genes in Ciona savignyi. Gene 705: 142–148. doi:10.1016/j.gene.2019.04.056

Ruiz GM, Carlton JT. 2003. Invasion vectors: a conceptual framework for management. In: Ruiz GM, Carlton JT (eds) Invasive species: vectors and management strategies. Island Press, Washington, pp 459–504.

Sadek J, Omer A, Hall D, Ashour K, Gallouzi IE. 2019. Alternative polyadenylation and the stress response. WIREs RNA 10(5): e1540. doi:10.1002/wrna.1540

Salisbury SJ, Delgado ML, Dalziel AC. 2021. Alternative splicing: An overlooked mechanism contributing to local adaptation? Mol Ecol 30(20): 4951–4954. doi:10.1111/mec.16177

Sandoval-Castillo J, Gates K, Brauer CJ, Smith S, Bernatchez L, Beheregaray LB. 2020. Adaptation of plasticity to projected maximum temperatures and across climatically defined bioregions. P Natl Acad Sci USA 117(29): 17112–17121. doi:10.1073/pnas.1921124117

Santi F, Riesch R, Baier J, Grote M, Hornung S, Jungling H, Plath M, Jourdan J. 2020. A century later: Adaptive plasticity and rapid evolution contribute to geographic variation in invasive mosquitofish. Sci Total Environ 726: 137908. doi:10.1016/j.scitotenv.2020.137908

Schaefke B, Sun W, Li YS, Fang L, Chen W. 2018. The evolution of posttranscriptional regulation. WIRES RNA e1485. doi:10.1002/wrna.1485

Shalgi R, Hurt JA, Lindquist S, Burge CB. 2014. Widespread inhibition of posttranscriptional splicing shapes the cellular transcriptome following heat shock. Cell Rep 7(5): 1362–1370. doi:10.1016/j.celrep.2014.04.044

Shen S, Park JW, Lu ZX, Lin L, Henry MD, Wu YN, Zhou Q, Xing Y. 2014. rMATS: robust and flexible detection of differential alternative splicing from replicate RNA-Seq data. P Natl Acad Sci USA 111(51): E5593–5601. doi:10.1073/pnas.1419161111

Si Y, Wen H, Li Y, He F, Li J, Li S, He H. 2018. Liver transcriptome analysis reveals extensive transcriptional plasticity during acclimation to low salinity in Cynoglossus semilaevis. BMC Genomics 19(1): 464. doi:10.1186/s12864-018-4825-4

Singh P, Borger C, More H, Sturmbauer C. 2017. The Role of Alternative Splicing and Differential Gene Expression in Cichlid Adaptive Radiation. Genome Biol Evol 9(10): 2764–2781. doi:10.1093/gbe/evx204

Smith K, Cahill P, Fidler A. 2010. First record of the solitary ascidian Ciona savignyi Herdman, 1882 in the Southern Hemisphere. Aquat Invasions 5(4): 363–368. doi:10.3391/ai.2010.5.4.05

Sokolov EP, Sokolova IM. 2019. Compatible osmolytes modulate mitochondrial function in a marine osmoconformer Crassostrea gigas (Thunberg, 1793). Mitochondrion 45: 29–37. doi:10.1016/j.mito.2018.02.002

Su H, Ma D, Zhu H, Liu Z, Gao F. 2020. Transcriptomic response to three osmotic stresses in gills of hybrid tilapia (Oreochromis mossambicus female x O. urolepis hornorum male). BMC Genomics 21(1): 110. doi:10.1186/s12864-020-6512-5

Subramanian A, Tamayo P, Mootha VK, Mukherjee S, Ebert BL, Gillette MA, Paulovich A, Pomeroy SL, Golub TR, Lander ES, et al. 2005. Gene set enrichment analysis: a knowledge-based approach for interpreting genome-wide expression profiles. P Natl Acad Sci USA 102(43): 15545–15550. doi:10.1073/pnas.0506580102

Suresh S, Crease TJ, Cristescu ME, Chain FJJ. 2020. Alternative splicing is highly variable among Daphnia pulex lineages in response to acute copper exposure. BMC Genomics 21(1): 433. doi:10.1186/s12864-020-06831-4

Tian B, Manley JL. 2017. Alternative polyadenylation of mRNA precursors. Nat Rev Mol Cell Bio 18(1): 18–30. doi:10.1038/nrm.2016.116

Tian Y, Wen H, Qi X, Zhang X, Sun Y, Li J, He F, Zhang M, Zhang K, Yang W, et al. 2020. Alternative splicing (AS) mechanism plays important roles in response to different salinity environments in spotted sea bass. Int J Biol Macromol 155: 50–60. doi:10.1016/j.ijbiomac.2020.03.178

Verta JP, Jacobs A. 2022. The role of alternative splicing in adaptation and evolution. Trends Ecol Evol 37: 299–308. doi:10.1016/j.tree.2021.11.010

Wiens JJ. 2016. Climate-Related Local Extinctions Are Already Widespread among Plant and Animal Species. PLoS Biol 14(12): e2001104. doi:10.1371/journal.pbio.2001104

Wright RM, Aglyamova GV, Meyer E, Matz MV. 2015. Gene expression associated with white syndromes in a reef building coral, Acropora hyacinthus. BMC Genomics 16: 371. doi:10.1186/s12864-015-1540-2

Yancey PH. 2005. Organic osmolytes as compatible, metabolic and counteracting cytoprotectants in high osmolarity and other stresses. J Exp Biol 208(15): 2819–2830. doi: 10.1242/jeb.01730

Ye C, Long Y, Ji G, Li QQ, Wu X. 2018. APAtrap: identification and quantification of alternative polyadenylation sites from RNA-seq data. Bioinformatics 34(11): 1841–1849. doi:10.1093/bioinformatics/bty029

Ye C, Zhou Q, Wu X, Ji G, Li QQ. 2019. Genome-wide alternative polyadenylation dynamics in response to biotic and abiotic stresses in rice. Ecotox Environ Safet 183: 109485. doi:10.1016/j.ecoenv.2019.109485

Zhan A, Briski E, Bock DG, Ghabooli S, MacIsaac HJ. 2015. Ascidians as models for studying invasion success. Mar Biol 162(12): 2449–2470. doi:10.1007/s00227-015-2734-5

Zhan A, MacIsaac, HJ, Cristescu, ME. 2010. Invasion genetics of the Ciona intestinalis species complex: from regional endemism to global homogeneity. Mol Ecol 19(21): 4678–4694. doi: 10.1111/j.1365-294X.2010.04837.x

Zhao X, Yu H, Kong L, Li Q. 2016. Gene co-expression network analysis reveals the correlation patterns among genes in euryhaline adaptation of Crassostrea gigas. Mar Biotechnol 18(5): 535–544. doi:10.1007/s10126-016-9715-7

Zheng D, Wang R, Ding Q, Wang T, Xie B, Wei L, Zhong Z, Tian B. 2018. Cellular stress alters 3’UTR landscape through alternative polyadenylation and isoform-specific degradation. Nat Commun 9(1): 2268. doi:10.1038/s41467-018-04730-7

